# Wildfire ash impacts to photosynthesis and growth of marine phytoplankton cultures

**DOI:** 10.1101/2025.01.10.632499

**Authors:** Kyle S. Van Houtan, John Lambert, Anthony A. Provatas, Dillon J. Van Houtan, Celia M. Smith

## Abstract

Climate change includes increasing surface temperatures as well as extreme events—heatwaves, storms, floods, and droughts. In 2020, these factors produced a record 10,000 wildfires that burned 18,000 km^2^ in California USA. Air pollution, including airborne ash, from these fires was a widespread human health hazard. While the ecological effects of wildfires have been extensively documented in terrestrial and freshwater systems, impacts on ocean and coastal ecosystems are largely unexplored. Here, we describe the physical and chemical properties of ash from the CZU Lightning Complex fire and experimentally test its effects on the photosynthesis and growth of four unicellular marine phytoplankton. Sieved air-fall ash was primarily composed of particles 250-500 μm and contained ∼1 ‰ of Fe, Mn, and Ba. Diagnostic indices of polycyclic aromatic hydrocarbons indicated the air-fall ash originated from combusted wood, and the total concentration of the EPA 16 high-priority PAHs exceeded 2.7 ppm. Pulse Amplitude Modulation fluorometry documented various declines in the photosynthetic efficiency of Isochrysis and Dunaliella cultures dosed with ash, and the bulk cellular growth of these cultures was inhibited. While our study demonstrated the impacts of wildfire ash on marine producers, the precise mechanisms are unclear. We provide recommendations for how future studies may further resolve the impacts of ash on phytoplankton productivity, community diversity, and trophic transfer of toxins and describe the long-term impacts of wildfires on coastal marine ecosystems.

## INTRODUCTION

Ocean phytoplankton are the foundation of marine food webs and determine ecosystem dynamics, biodiversity patterns, and global biogeochemical cycles (Falkowski 1994; Gagné et al. 2020; Litchman et al. 2015; Ridgwell and Zeebe 2005). It is a significant concern, therefore, that marine phytoplankton have declined broadly over the last century (Boyce et al. 2010). Given the bottom-up forcing of temperature and ocean currents to the key physiology and processes of marine phytoplankton, global climate change is considered the primary cause of these observed changes (Beardall and Raven 2004; Winder and Sommer 2012). In contrast to these known climatic drivers, the additional influence of anthropogenic pollution to marine phytoplankton is not well established.

Even when benchmarked by near-historical baselines, catastrophic impacts arising from extreme climate change are increasingly common (Grillakis 2019; Tanaka and Van Houtan 2022; Tripathy et al. 2023; Walsh et al. 2016). Perhaps it is therefore unsurprising that terrestrial wildfires driven by climatic drought and warming have become more frequent and widespread, and that our world is dangerously more fire prone (Cunningham et al. 2024). The collective socioeconomic and public health impacts from wildfires are staggering. Today, wildfires constitute a significant proportion of the total global harmful particulate air pollution and incur in excess of $100 billion USD damages annually (Black et al. 2017; Wang et al. 2021). And while the impacts of wildfires and their associated pollution has been documented in terrestrial and aquatic ecosystems (Earl and Blinn 2003; Gajendiran et al. 2024; Klose et al. 2015), we do not presently understand their potential impact in marine systems—either currently or in future climate scenarios. What is the impact of wildfires for the ocean and coastal ecosystems?

The CZU Lightning Complex (“CZU”) fire burned 350 km^2^ of coastal California from August to September 2020. Before the fire, land cover within the CZU perimeter consisted primarily of evergreen forest, mixed forest and chaparral ecosystems set within a steep mountain terrain and containing multiple ocean-connected watersheds (Luft et al. 2022; Potter 2023). Due the dry conditions and intensity of the CZU fire, its ecological damage was severe, with > 75% tree mortality after two years (Potter 2023). Beyond the immediate and acute threats of the destruction of natural and human infrastructures, the CZU fire posed additional pollution risks. Smoke and ash from combusted vegetation reduced air quality to hazardous levels—i.e., the concentration of particulate air matter < 2.5 µm (“PM_2.5_”) exceeded 35.5 µg m^3^ (EPA 2024)—for weeks (Figures 1, 2a). While impacts from the CZU fire are still being documented, an understanding of their impacts to coastal and marine systems is largely unexplored (Kramer et al. 2020; Takesue et al. 2023).

**Figure 1.**
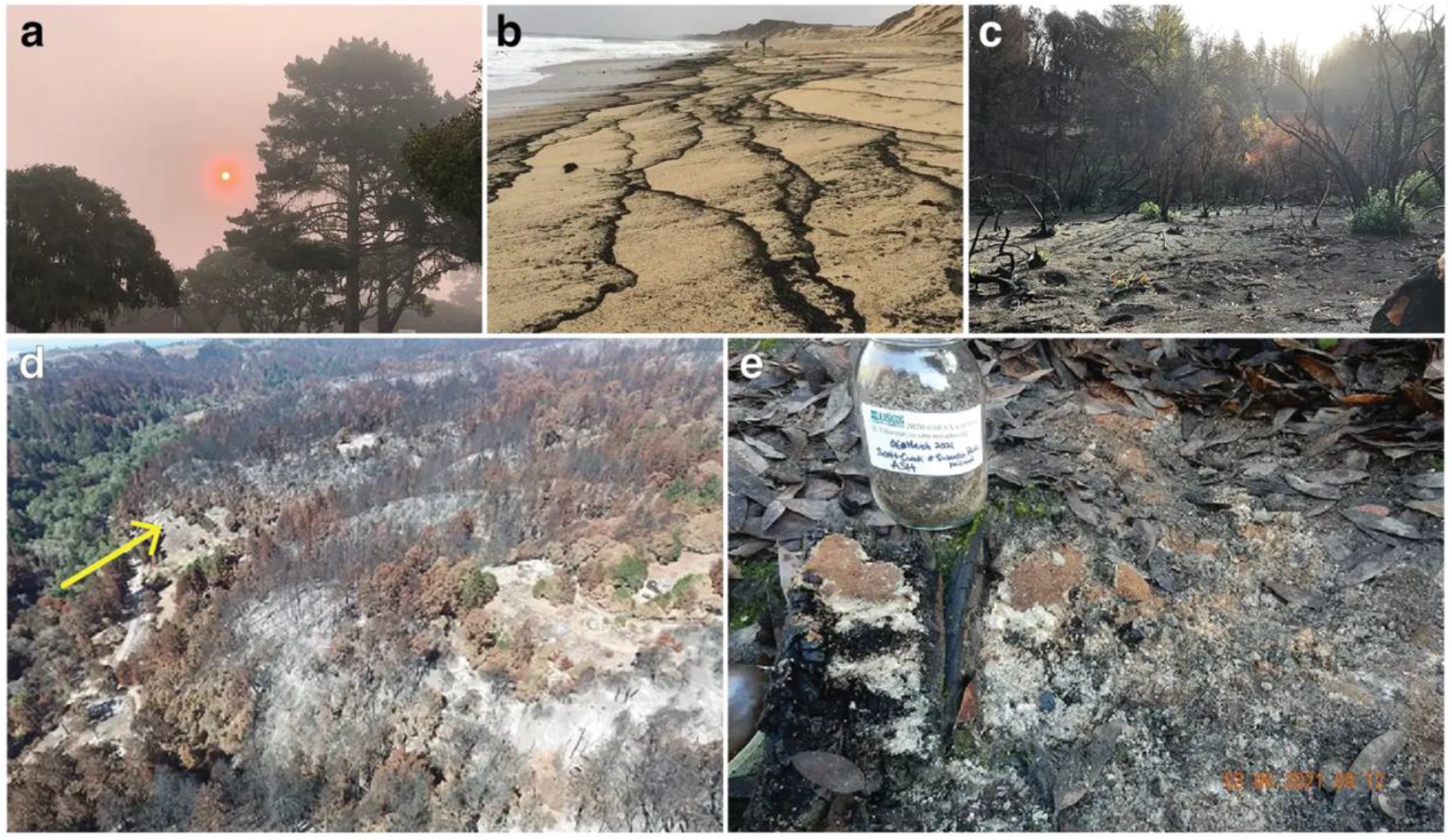
Wildfire ash from the CZU Lightning Complex fire. Smoke and ash from the CZU fire caused severe air pollution in central California during Aug-Sep 2020. (a) Air pollution from the CZU in Pacific Grove, CA on 8/21/20 when air-fall ash was collected (Figure 1) and community air quality sensors sustained PM_2.5_ levels > 200 µg m^−3^. (b) Air-fall ash from the CZU fire accumulated in coastal environments (charcoal contours) with unknown ecosystem impacts. Image taken 8/27/20 at Del Monte Beach in Sand City (36.620° N, 121.849° W, image: J. O’Sullivan). (c) CZU_1_ site of ground-collected ash near the San Vicente Redwoods natural area (37.105° N, 122.144° W), taken 12/12/20. (d) Aerial image of the CZU_2_ site in the Scott Creek watershed, near Swanton Road (37.073° N, 122.230° W) taken 3/5/21 (image: K. Kittleson/County of Santa Cruz). Yellow arrow denotes collection site. (e) CZU_3_ site along the bank of Scott Creek at Swanton Pacific Railroad (37.062° N, 122.229° W), taken 3/6/21 (image: R. Takesue/USGS). All images used with permission. Images from the authors unless otherwise noted.

**Figure 2.**
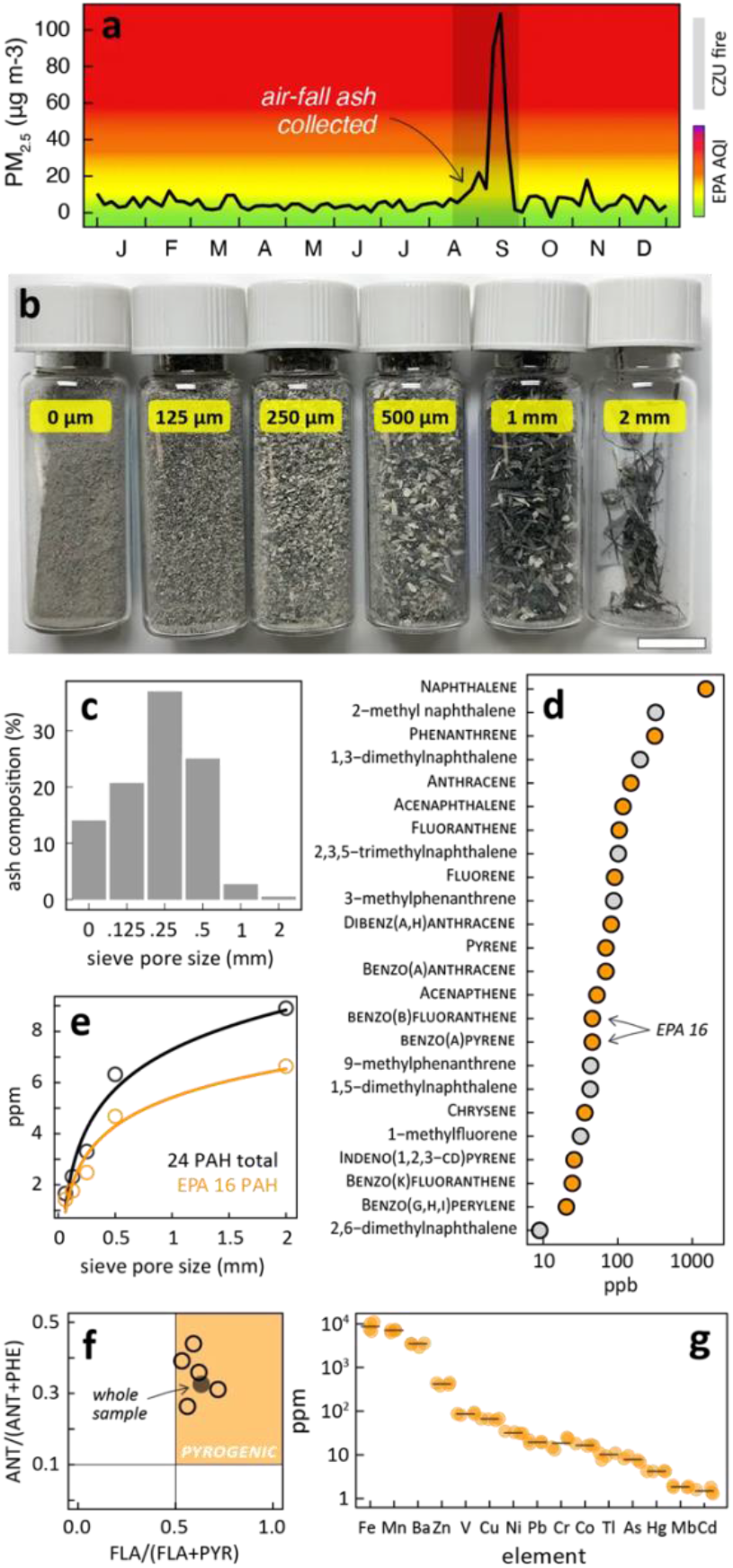
Physical and chemical properties of air-fall ash from the 2020 CZU fire. (a) *In situ* sensors in Pacific Grove, California indicate air pollution reached hazardous levels for human health during the CZU fire (shaded box). Solid line is the 24-hour running mean. Color ramp is the US Environmental Protection Agency Air Quality Index (EPA 2024). Sieved air-fall ash contained (b) partially combusted woody materials and conifer needles and was primarily composed of (c) particles between 250-500 μm. Air-fall ash had (d) elevated polycyclic aromatic hydrocarbon (PAHs) levels and included the 16 high priority PAHs (orange-filled circles, labels in SMALL CAPS) designated by the US Environmental Protection Agency (EPA). (e) PAH concentrations increased with particle size, and (f) diagnostic PAH indices of anthracene (ANT) and phenanthrene (PHE), and fluoranthene (FLA) and pyrene (PYR) indicate that the air-fall ash originated from combusted wood. Open circles are size-fractionated ash samples, filled grey circle is whole sample. (g) Fe, Mn, and Ba concentrations in whole sample air-fall ash exceeded 1‰, orange circles are bootstrap samples.

In this study, we developed *ex situ* aquaculture experiments to examine how wildfire might affect marine phytoplankton. First, we collected ash samples and examined their stable isotope, heavy metal (“HM”), and polycyclic aromatic hydrocarbon (“PAH”) content. Next, we assessed how ash dosages influenced the growth and photosynthetic efficiency of controlled monocultures of Haptophyta and Chlorophyta plankton. We then used random forest (“RF”) algorithms to build machine learning models to rank and evaluate a suite of potential model predictors. These models help assess the various potential inhibiting (HM and PAH toxicity, light shading) and fertilizing (Fe, C, N) impacts from wildfire ash to phytoplankton, as well as time and taxonomic effects.

## METHODS

### Air quality monitoring, Wildfire ash collection, and diagnostic analysis

Purple Air instruments documented community air quality for the 2020 calendar year before, during, and after the CZU fire. These open-access, low-cost commercial platforms effectively describe air pollution in the absence of regulatory-grade instrumentation (Feenstra et al. 2019; Jaffe et al. 2022). We obtained particulate matter below 2.5 µm (PM_2.5_) data from 3 separate Purple Air stations in Pacific Grove, California. For each station, we analyzed the primary data series (https://bit.ly/3GsdpMw) from both A and B channels where each sensor (Plantower, no. PMS5003) detects PM_2.5_ concentrations ≤ 500 μg m^−3^. Following best practices, PM_2.5_ values are the average of the “PM2.5_CF1_ug/m3” series from all channels (at each 2-minute intervals) that we standardized with a correction algorithm (Barkjohn et al. 2021). To the resulting series of all stations, we fit a locally-weighted regression (“LOESS”, Cleveland and Devlin 1988) with a span of 0.001, for a ∼24-hour running mean. Figure S1 provides additional details on the sensors and data.

We sampled ash from air-fall deposition, burn site collections, and experimental combustion (see Figure 1). At the outset of the CZU fire, from 19-21 August 2020, we passively collected air-fall ash in pre-cleaned glassware situated on the roof of the Monterey Bay Aquarium (36.6181° N, 121.9020° W). After the fires ended, research partners collected ash from CZU burn scars. In all field sites, collectors scraped superficial ash from undisturbed ground surfaces with pre-cleaned instruments and stored ash in cleaned containers. National Oceanic and Atmospheric Administration staff collected CZU_0_ samples on 11/5/20 near Big Creek Road (37.0764° N, 122.2194° W). CZU_1_ samples were collected on 12/12/20 near the San Vicente Redwoods natural area (37.1048° N, 122.1435° W). County of Santa Cruz staff collected CZU_2_ samples on 3/5/21 in the Scott Creek watershed (37.0729° N, 122.2306° W). U.S. Geological Service staff collected CZU_3_ samples on 3/6/21 (see Takesue et al. 2023) near the Swanton Pacific Railroad (37.0621° N, 122.2286° W). We also generated ash experimentally, by combusting foliage and branches from 7 tree species representative of local taxa (collected in Pacific Grove) in a pre-cleaned outdoor wood oven. Tables S1-S2 provide details on all ash samples.

We sieved and analyzed ash samples for their material composition. As CZU_1_ samples were collected during heavy rains, we desiccated CZU_1_ ash at 60°C overnight in a drying oven (OVG-400-27-120 Gravity Convection, Cole-Parmer). US standard (ASTM E-11) stainless steel sieves (8” x 2”, Fisherbrand™) separated dried ash particles on 2mm, 1mm, 500µm, 250µm, and 125µm mesh surfaces, and smaller particles. We kept size fractionated ash in sterile glassware, sending samples in 4 mL glass screw-top vials (Thermo Scientific, 13 mm ID) for HM screening at the California Animal Health and Food Safety Laboratory (e.g., Gagné et al. 2019), and for stable isotope analysis at the Georgia Center for Applied Isotope Studies (e.g., Van Houtan et al. 2023).

Following Provatas et al. (2019) and Chiesa et al. (2018), we quantified PAH content using gas chromatography with tandem mass spectrometry (“GC-MS/MS”) at the University of Connecticut, Center for Environmental Sciences and Engineering. We inserted a 0.2 g aliquot of ash to a 2 mL centrifuge tube, added 1 mL of CH_3_CN, and 0.3 g of Quick, Easy, Cheap, Effective, Rugged, and Safe extraction (“QuEChERS”, MgSO4/PSA). We vigorously vortexed samples for 15 minutes and centrifuged for 10 minutes at 14,000 RPM. From each sample, we transferred a 190 µL aliquot of the CH_3_CN extract into an analytical vial and spiked it with crysene-d12 internal standard. A Waters Quattro Micro GC-MS/MS system controlled by MassLynx software (v4.2) analyzed all samples, including quality controls. GC-MS/MS used a Restek Rxi-5Sil MS (30 m × 0.25 mm id × 0.25 μm) with the ensuing temperature routine: 90°C for 0.5 min, then 9°C min^−1^ ramp to 171°C, 2°C min^−1^ ramp to 176°C, 3°C min^−1^ ramp to 222°C, 2°C min^−1^ ramp to 242°C, 10°C min^−1^ ramp to 280°C, and hold at 280°C for 10 min. We used He carrier gas at a constant flow 1.0 mL min^−1^, Ar collision gas (3.0 × 10^−3^ mbar), inlet temperature 280°C, injection volume 1 μL (splitless), MS transfer line temperature at 280°C, ion source temperature at 250°C, with an electron impact ionization MS mode at 70 eV. QuanLynx software analyzed analyte outputs. Diagnostic PAH indices (e.g., Takesue et al. 2023) confirmed air-fall ash origin.

### Phytoplankton cultures and experimental design

Online Supplementary Material describes the aquaculture facilities and preparations for the phytoplankton cultures. For pilot and experimental cultures, carboy containers were spaced evenly on shelves in a naturally sunlit, well-ventilated, and dedicated culture room. Each shelving system was outfitted with two covered 4’ fluorescent light fixtures (Lithonia Lighting, Clear diffuser lens, 2 tubes fixture^−1^), each equipped with 13W LED tubes (Green Creative Titanium, 4000K), anchored ∼45 cm above the array. Light assemblies operated continually to maximize culture production. We rotated container shelf orientations and changed positions daily to avoid potential microclimate effects. Culture temperatures were ambient, unregulated, and did not vary appreciably over the experiment.

During a pilot testing phase, we added 10 mL liquid Guillard’s F/2 seawater enrichment medium (Micro Algae Grow™, Florida Aqua Farms Inc.) to each culture (Guillard 1975; Guillard and Ryther 1962). Commercial aquaculture suppliers (Florida Aqua Farms Inc., AlgaGen LLC) provided stocks of *Isochrysis galbana* (Haptophyta), *Dunaliella tertiolecta* (Chlorophyta), *Tetraselmis chuii* (Chlorophyta), and *Rhodomonas lens* (Cryptophyta). We separately cultured phytoplankton in large columns for the maintenance and sustainability of bivalve populations (Creswell 2010) that support public-facing exhibits at the Monterey Bay Aquarium. We inoculated 8 carboys with phytoplankton cultures (2 carboy chambers × 4 species), removing 2 L of dense algae inoculant from the healthy culture source and adding to distinct carboys. Over a period of 28 days, we measured their photosynthetic efficiency (see below) to calibrate the length of the experiments and determine the best experimental models.

For experiment cultures, we added 5 mL of Guillard’s medium to avoid masking ash impacts. From perceived robustness, taxonomic representation, and logistical constraints, we inoculated 3 carboys each with *I. galbana* and *D. tertiolecta*. For a positive control, 1 container of each species received no additional treatments. For the experimental treatments, on day 4 after cultures were inoculated, we dosed 2 containers of each species with ash. One container received 1.23 g L^−1^ of ash treatment 1 (“ash 1”), the other a similar amount of ash treatment 2 (“ash 2”). Ash 1 was composed of 3.3% air-fall ash, 88.5% CZU_0_ ash, and 8.3% CZU_1_ ash. Ash 2 is 55.1% CZU_1_ ash and 44.9% experimentally generated ash. We desired to use air-fall ash in the experiments, but these samples were consumed by diagnostic laboratory analyses. For the negative controls, we prepared 2 carboys with seawater, added no phytoplankton, and immediately dosed them with 1.23 g L^−1^ of ash. This resulted in 10 total experimental containers: 2 species × 2 ash treatments, 2 positive controls (phytoplankton without ash), 2 negative controls (ash without phytoplankton), plus 2 more blank containers of sterilized seawater. Table S3 provides additional details on the experimental treatments and controls.

### Assessing culture photosynthesis and growth

We monitored phytoplankton cultures for cell growth and photosynthesis. For growth, we modified methods previously used in monitoring microplastic and eDNA (Choy et al. 2019; Searcy et al. 2022). Here, we assembled a 47mm diameter glass filtration set (All-Glass Filter Holder kit, Millipore Sigma) consisting of a 300 mL funnel, anodized Al clamp, glass frit membrane support, and 1 L receiver flask attached to a vacuum (Cast N’ Vac Vacuum 1000, Buehler™). Initially, we used 0.3 µm glass microfiber prefilters (A/D Glass Fiber 47mm, PALL Corporation). However, as these materials had supply chain challenges, slowed the process considerably, and as the minimum cell volume of our phytoplankton is > 40 µm^3^ (Creswell 2010), we switched to 1 µm filters (GF/F 47mm, Whatman™).

We filtered culture extracts and transferred used filters to covered borosilicate glass petri dishes (reusable 50mm, PYREX™), kept them at 60°C overnight in a drying oven, and weighed dried filters on a microbalance readable to 0.0001 g. We repeated this routine for extracts from blank chambers (4 measurements on 2 days), and negative controls (8 measurements on 4 days). To isolate and calculate phytoplankton growth in the positive controls, we subtracted mass of their pre-weighed empty filters and the mean mass of filtered dried seawater extracts from the mass of dried positive control filters. To isolate and calculate phytoplankton growth in the experimental treatments, we subtracted the mass of their pre-weighed empty filters, and the mean mass of the filtered negative control extracts from the mass of dried experimental filters. For all phytoplankton cultures, we performed daily filtrations, extracting 50-150 mL of solution from each chamber, expressing growth as mg of phytoplankton dry mass per mL of culture solution.

A portable photosynthesis yield analyzer (MINI-PAM-II/B, Walz Corporation) operated by WinControl software (v3.3) measured Pulse Amplitude Modulation (“PAM”) fluorometry (Anonymous 2024) of phytoplankton cultures. PAM calculated the effective photochemical quantum yield of Photosystem II (“PS II”) in phytoplankton cultures using:

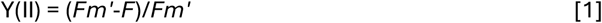

where *F* is the recorded fluorescence yield of cultures before illumination and *Fm’* is the observed maximum fluorescence yield of illuminated cultures (Klughammer and Schreiber 2008). The MINI-PAM analyzer generated experimental light through LED fiberoptics, using the saturation pulse method. The parameter Y(II) is a metric of photosynthetic efficiency as it captures “the fraction of energy that is photochemically converted in PS II” and as its complement, 1-Y(II), represents the fraction of photochemical energy PS II loses (Klughammer and Schreiber 2008). Consequently, Y(II) is a key index for assessing the performance/stress to photosynthesis across taxa (Amato et al. 2018; Murchie and Lawson 2013; Padilla-Gamiño et al. 2019).

Considering our experimental aquaculture setting, and following manufacturer advice (Anonymous 2024), we operated the PAM analyzer by placing the tip of the fiberoptics adapter directly flush, 90° to surface of pre-cleaned culture container surfaces. PAM illuminated cultures after 20 minutes of dark adaptation. We dark adapted phytoplankton cultures by covering carboys with dark, thick, and insulated winter outerwear. We fitted covers tightly around each container, allowing the air inlet and outlet tubes to vent, and ensuring carboys remained completely shrouded. We took PAM measurements by inserting the fiberoptics adapter through a small seam in the covers, and assessed the rigor of Y(II) measurement by inspecting the fluorescence kinetics curves for a canonical asymptotic form (Anonymous 2024). We recorded 2-3 Y(II) measurements for each container at one observation period daily, initiating measurements on the day of culture inoculation and continuing measurements daily.

### MODELING DRIVERS OF PHOTOSYNTHETIC EFFICIENCY

To understand culture performance, RF algorithms fitted multiple non-linear regressions (Breiman 2001) to the Y(II) data with a suite of likely drivers. We previously applied RF in similar contexts to model the complex influences of likely variables to ecological processes (Becker et al. 2019; Becker et al. 2020; Gagné et al. 2018a; Gagné et al. 2018b; Nicholson et al. 2023; Nicholson et al. 2024). RF is an appealing modeling framework as it accommodates heteroskedasticity and nonlinearity, has built-in procedures to reduce bias and overfitting, incorporates both continuous and categorical predictors, and provides tools to visualize the joint influence of multiple predictors to dependent variables (Breiman 2001; Greenwell 2017; Van Houtan et al. 2024).

We performed all analyses in the R computing environment (version 4.2.2, RCoreTeam 2022) run in macOS Sonoma 14.6.1 using an 8-core M1 chip. The ‘dplyr’ and ‘tidyr’ libraries provided essential functions for wrangling, joining, and bootstrapping data (Wickham et al. 2023). From the measured Y(II) observations, we calculated the mean and standard deviation (“sd”), after grouping the data by day and carboy chamber (11 days × 2 species × 3 treatments = 66 groups). When Y(II) observations were equal (i.e., sd=0), we used the mean sd across groupings. The ‘replicate’ and ‘mutate’ functions performed parametric bootstrapping, generating 400 Y(II) values from each mean and sd, for a total of 26400 Y(II) values. The full suite of model predictors included 11 categorical and continuous variables. Three categorial variables were phytoplankton genus (*Dunaliella, Isochrysis*), treatment type (control, ash1, ash2), and ash exposed (yes, no). Eight continuous variables were culture age (0-12 days), time since ash exposure (0-8 days), ash dosage (0-28 g), ash C content (0-2.9 g), 16 EPA PAH (0-18 µg), total PAH (0-24 µg), ash Fe (0-147 mg), and 15 HM totals (excluding Fe, 0-117 mg). Added C, PAH, and HM were from ash doses, derived from the diagnostic results presented in Figures 2-3.

The ‘caret’ and ‘RandomForest’ libraries trained, tuned, and ran RF models (Kuhn et al. 2020; Liaw and Wiener 2002). We initially tuned and ran RF with all covariates (n=11) but reduced to a core suite (n=5) of independent variables representative of the full suite of predictors, without performance loss. The final 5 predictors were: genus, treatment type, culture age, Fe dosage, and total PAH dosage. RFs were trained using a resampling routine of 10-fold cross validation with 5 repetitions (Gareth et al. 2013), fixing the number of model trees (*ntree* = 2000), and tuning the random node splitting (*mtry*) for optimal performance (determined by RMSE). The ‘doParallel’ package enabled parallel processing (Weston and Calaway 2022). The ‘ggplot2’ and ‘pdp’ libraries provided data wrangling and visualization tools to aid model design and interpretation (Greenwell 2017; Wickham 2011).

## RESULTS

Smoke and ash from the CZU wildfire reached the Monterey Peninsula (Figure 1a-b) >50 km distance from the fire boundary, with unknown coastal impacts. In addition to air-fall deposition collected during active fires, we obtained ash from CZU burn scars (Figure 1c-e) after fires ceased. Purple Air stations provided 1,951,066 PM_2.5_ measurements over 1,703 sensor days. 24-hour running mean of these data indicated hazardous air quality (PM_2.5_ > 35.5 µg m^−3^) throughout September 2020, during the CZU fire.

Deposited air-fall ash contained partially combusted conifer needles (Figure 1b) and size fractionated air-fall ash was primarily composed of particles 250-500 µm in size (Figure 1c). Of the 24 PAHs quantified, the highest concentrations were primarily low molecular weight compounds the EPA has prioritized due to their acute toxicity, higher solubility and fast absorption rates (Geier et al. 2018). Only naphthalene exceeded a whole sample concentration > 1 ppm (1.52 ppm), while the whole sample had 3.6 ppm of all 24 PAHs, and 2.77 ppm for the 16 PAHs prioritized by the U.S. Environmental Protection Agency (Figure 1d). The total ash concentration of all 24 PAHs and the 16 EPA priority PAHs increased with particle size, following a logarithmic form (Figure 1e), suggesting thorough combustion volatilizes PAHs. Figure 1f shows 2 diagnostic PAH ratios that independently document air-fall ash arose from wood combustion (Takesue et al. 2023). The ratio of anthracene to anthracene + phenanthrene exceeded 0.1 in all size-fractionated samples, indicated a pyrogenic source. The ratio of fluoranthene to fluoranthene + pyrene exceeded 0.5 in all size fractionated samples, indicated wood combustion. HM screens documented air-fall ash was 96.6% of Fe (8.9‰), Mn (7.0‰), and Ba (3.4‰), with all other HM elements combined totaling < 0.7‰ (Figure 1g). Due to limited air-fall ash samples, we did not size fractionate HM content.

The size and chemical profile of CZU ash collected *in situ* and experimentally generated ash was similar to air-fall ash, with noted differences. Collected CZU ash (combined from all 4 locations) was primarily composed (52.2%) of particles 250-500 µm in size. However, the relative proportion of generated ash increased as particle size decreased, with the largest proportion (31.0%) in the smallest particle size (0-125 µm, Figure 3a). While naphthalene and phenanthrene were leading PAHs in collected and generated ash, some low molecular weight PAHs (e.g., the methylnaphthalene isomers) declined (Figure 3b). The accumulated PAH concentrations were lower than air-fall ash, with collected ash = 0.54 ppm (all 4 locations samples combined) and generated ash = 0.73 ppm for all 24 PAHs. For the CZU_1_ samples of collected ash, size-fractionated HM concentrations show a mid-domain peak, with 9/15 elements achieving maximum concentrations in the 250-500 μm particle class (Figure 3c). (Be is not plotted as its HM content is < 1 ppm.) Like air-fall ash (Figure 2g), Fe, Mn, and Ba were dominant, but total HM content was lower in both (collected = 7.9‰, generated = 10.7‰). Stable isotope analysis indicated the CZU_1_ ash was 10.1% C, 0.05% N (Figure 3e). Stable C isotopes of CZU_1_ ash peaked between -26 and -25 δ^13^C, but δ^15^N was not quantified (Figure 3f).

**Figure 3.**
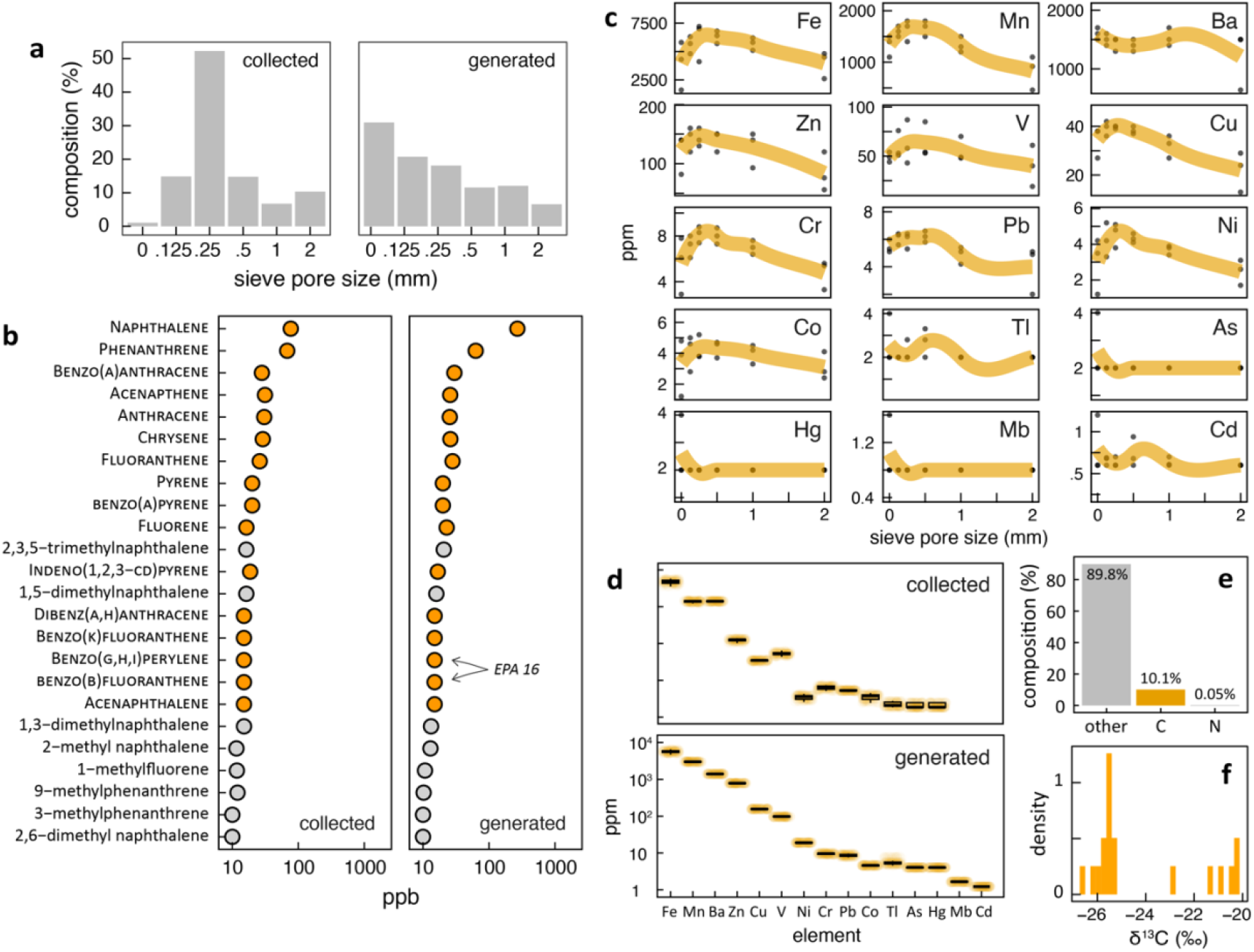
Physical and chemical properties of ground-collected wildfire ash and experimentally generated wood ash. (a) Ground-collected ash from CZU burn scars had a similar particle size distribution to air-fall ash (peak of 250-500 μm, see Figure 1), but experimentally generated ash was characteristically finer. (b) Collected and generated ash had lower PAH concentrations than air-fall ash. (c) Collected ash exhibited size-fractionated HM concentrations, with a peak at 250-500 μm sized particles for 9 of 15 elements tested. (d) Whole samples of collected and generated ash had comparable HM concentrations, similar to the relative composition of air-fall ash (Figure 1). Collected ash contained (e) 10.1% carbon, 0.05% nitrogen, and δ^13^C values (f) ranged from -27 to -20‰ (but mostly < -25‰). Low nitrogen concentrations inhibited δ^15^N quantification.

Phytoplankton were reared in 24-hour light conditions and dark adapted for 20 minutes before measuring Y(II) (Figure 4a-b). A 4-week trial of Haptophyta, Chlorophyta, and Cryptophyta cultures showed Y(II) increased rapidly from days 0-8, remained constant from days 8-12, and declined after (Figure 4c). Though we did not systematically sample all species throughout the period, the *Rhodomonas* culture had the highest measured Y(II) values (0.784, day 9). Solid line is an ensemble loess model for all 4 species. From this curve, we set our experiment period to 12 days.

**Figure 4.**
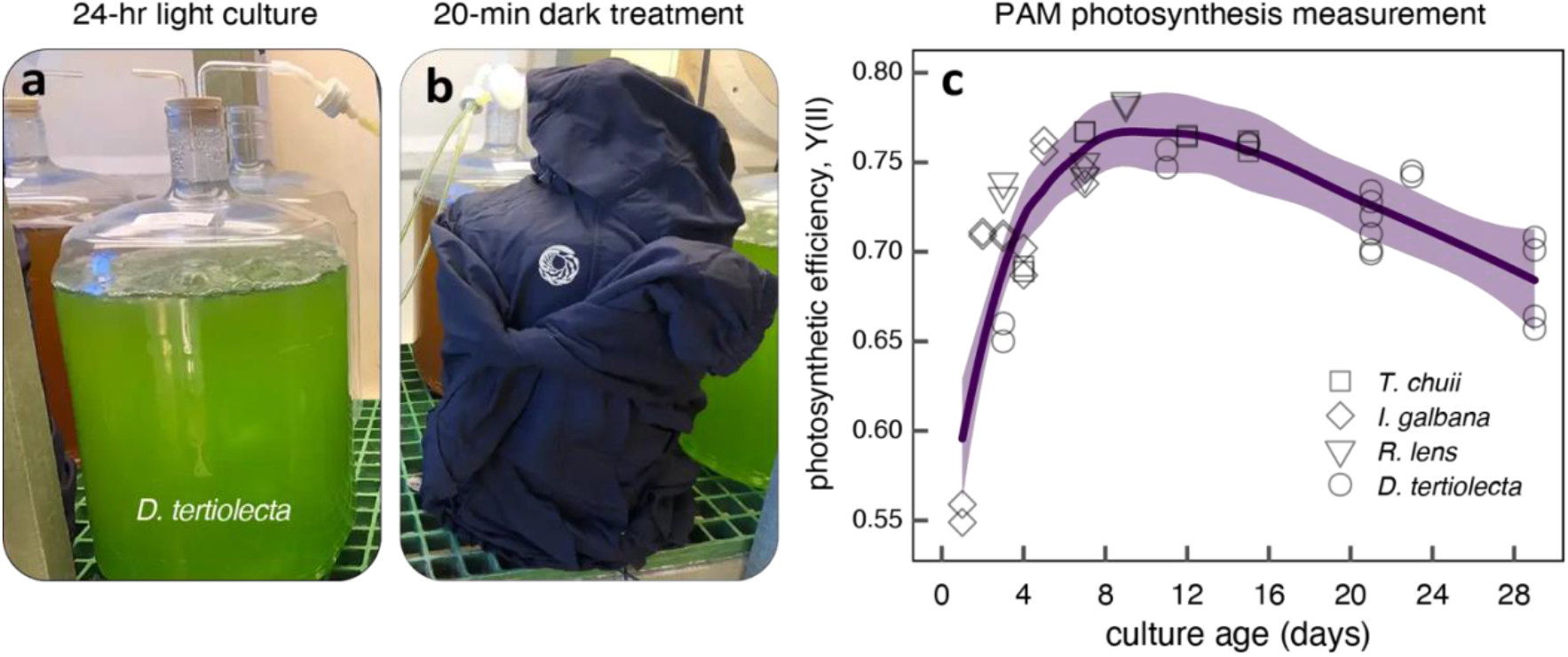
Idealized photosynthetic efficiency curves of cultured marine phytoplankton. We cultured four species of flagellate marine phytoplankton (*Isochrysis galbana, Dunaliella tertiolecta, Tetraselmis chuii*, and *Rhodomonas lens*) (a) continuously in 24-hour light conditions for one month to sustain exhibited bivalve populations at the Monterey Bay Aquarium. To assess phytoplankton photosystem operations, we (b) shrouded culture chambers for 15 minutes in dark conditions. While remaining dark, (c) Pulse Amplitude Modulation (PAM) fluorometry then captured the quantum yield of photochemical energy converted in photosystem II—known as photosynthetic efficiency, or Y(II). The ensemble trend of Y(II) for cultures over a typical one-month rearing cycle shows impacts to photosynthetic efficiency, suggesting negative density-dependence. Y(II) increased sharply (days 1-7), plateaued (days 8-12, with a peak median ensemble trend of 0.767 at day 9.5), and subsequently declined (days 13-29). These data calibrated the length of our experimental study to 12 days.

Figure 5a-b shows culture growth on dried filters from all three experimental phytoplankton treatments (no ash, ash 1, ash 2) as well as ash only treatments. A visual inspection of filtered culture extracts shows the gradual saturation of green (*D. tertiolecta*) and brown (*I. galbana*) cells as cultures mature—except for the ash 2 treatment *D. tertiolecta* which is inhibited. Filtered culture growth indicates the controls of both *D. tertiolecta* and *I. galbana* cultures outperform both ash treatments, with *D. tertiolecta* having more absolute growth (Figure 5c). *I. galbana* cultures attain higher Y(II) values (max = 0.76) than *D. tertiolecta* (max = 0.72), with both species achieving peak values after 8 days (Figure 5d). The Y(II) of ash 2 treatment of *D. tertiolecta* obviously drops after the ash is added. Negative controls had a median observed Y(II) value of 0.05 (*n* = 13).

**Figure 5.**
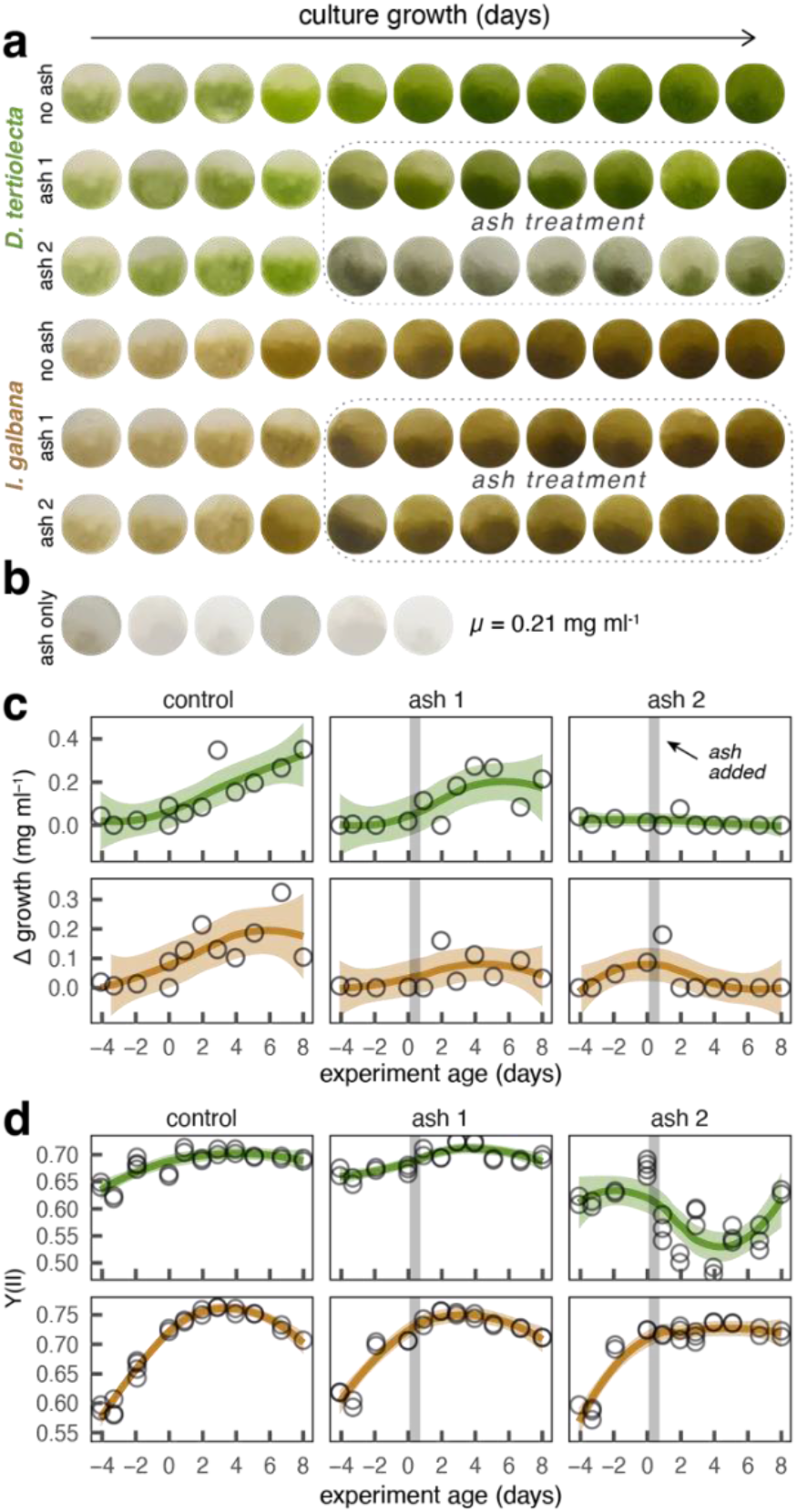
Experimental impacts of wildfire ash to the growth and photosynthesis of Haptophyta and Chlorophyta cultures. Vacuum-filtered culture extracts on glass microfibers captured (a) growth of *D. tertiolecta* and *I. galbana* cells under experimental exposures to combusted wood ash. Measurements were not taken on study days 3, 10 (experiment days -1, 6). Measured growth of ash-dosed cultures deducted the density of (b) seawater dissolved ash (*µ* = 0.21 mg ml^−1^). (c) Control groups (no ash) for both species show gradual colony growth throughout the study. For both species, ash 2 reduces growth more than ash 1, and *I. galbana* displays smaller differences between control and experimental treatments than *D. tertiolecta*, suggesting the former is more resistant, while the latter indicates recovery and resilience. Culture mass in (c) is expressed relative to the minimum daily value of each species group (Δ mg ml^−1^). Like the idealized Y(II) curves (Figure 4c), control cultures show (d) negative density-dependence in Y(II). Here, *I. galbana* has a higher variability of observed Y(II) values (median range: 0.574-0.761) than *D. tertiolecta* (range: 0.635-0.702). As in growth (c), ash impacts to (d) Y(II) in *D. tertiolecta* are more pronounced, with ash 2 treatments reducing Y(II) in both species. (c-d) Solid lines are loess regressions, shaded regions are standard errors, and vertical gray lines indicate date of ash application.

Pairwise comparisons of potential drivers display the relationships between experimental factors and Y(II). Figure 6a-b isolates the impacts of time and species on Y(II), showing the saturation at approximately day 8 and the elevated Y(II) of *I. galbana*. The impact of bulk culture treatment, Fe from ash, and PAH from ash show mixed effects (Figure 6c-e). While the control and ash 1 treatments have similar Y(II) values (median = 0.70) the ash 2 treatment is lower (median = 0.63, Figure 6d). Fe and PAH content (Figures 2-3) of ash treatments produces higher and lower Y(II) responses (Figure 6c,e). Random Forest regressions have a high explanatory value (*R*^2^ = 0.97), ranking culture age, taxon, and Fe as the top model predictors (Figure 6f). Partial dependency plots (Figure 6g-j) display multiple variable interactions on centered model predictions of Y(II), or *y^*. Despite the previously established patterns of culture age, taxon (Figure 6g), and ash treatment (Figure 6i), Figure 6h shows a slight fertilizing effect from added Fe.

**Figure 6.**
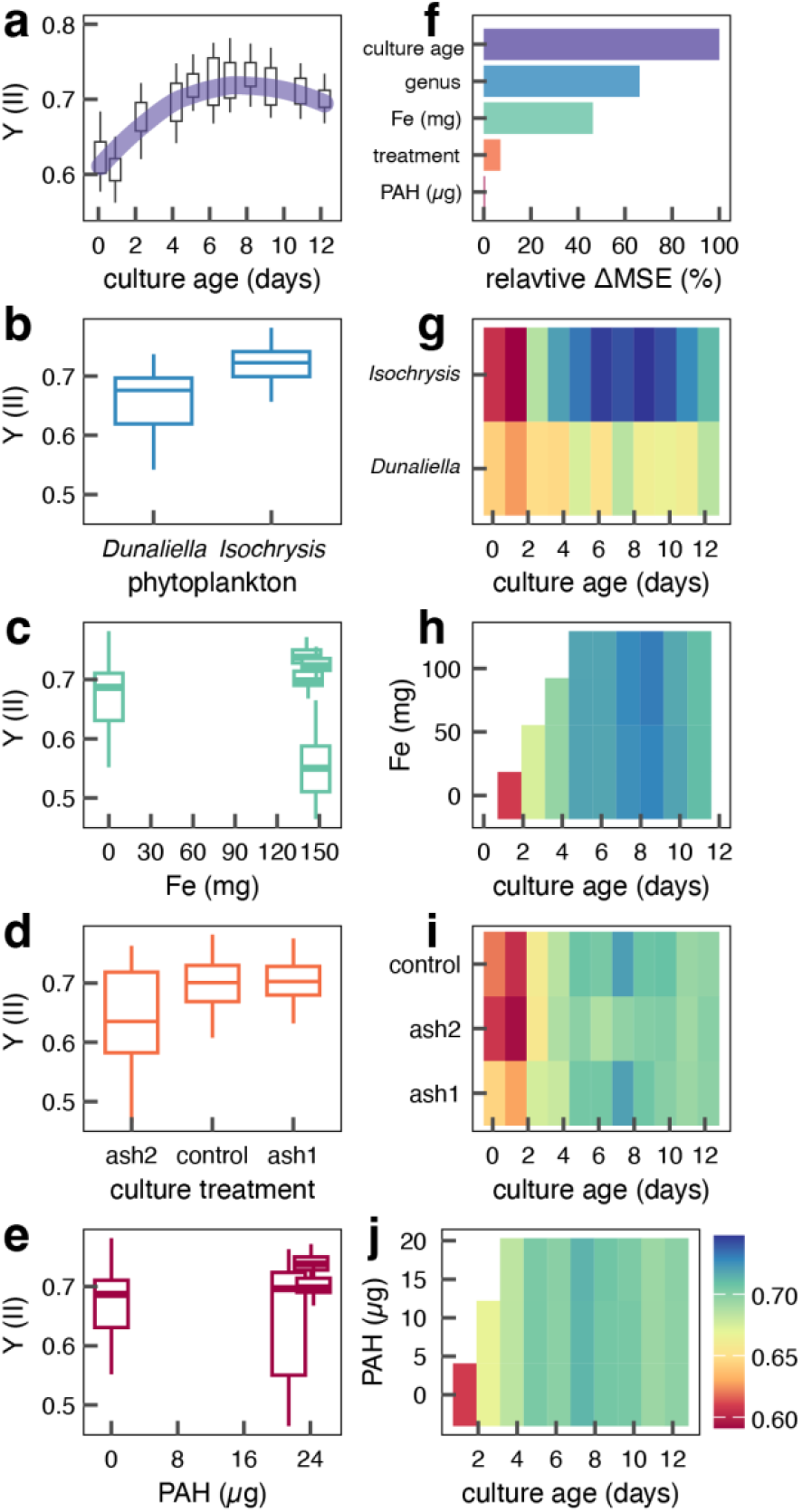
Culture age, phytoplankton taxon, and Fe are the top influences on Y(II), the photosynthetic efficiency of cultures. (a-e) Raw, pairwise relationships between the 5 model predictors and Y(II). (a) Combined data shows that cultures mimicked the idealized growth curve of the testing phase (Figure 4c) and summarize the raw experimental observations (Figure 5d). (b) *Isochrysis* outperforms *Dunaliella* cultures in Y(II), (c) Fe has mixed effects, (d) ash2 exhibits a clear decline in Y(II), where (e) PAH additions also produce mixed results. (f) Pre-trained and optimized RF models (*ntree* = 2000, *mtry* = 5) rank culture age, genus, and Fe as the top model drivers (*R*^2^ = 0.97), with bar symbology corresponding to the colors in (a-e). (g-j) Partial dependency plots display final model outputs, with model predictions (centered *y^*) of Y(II). Note that ash was added to experimental carboys on day 4, with (h) a slight increase in Y(II) on days 7-9 with Fe doses.

## DISCUSSION

While previous research has investigated the impacts of wildfire ash in terrestrial aquatic ecosystems, this is the first known study to assess its impacts on marine phytoplankton in an experimental setting. Air-fall, ground-collected, and experimentally generated ash all had a pyrogenic origin (Figure 1, 2f), yet the particle and toxicity profiles of these samples varied (Figures 2-3). Ash pollutants were notably size fractionated (Figures 2c, 3e), thus affecting their potential organismal impacts (e.g., Choy et al. 2019). Visual observations (Figure 5a) did not fully capture the changes to phytoplankton cultures from added ash, measured more precisely with absolute cellular growth (Figure 5c) and photosynthetic efficiency (Figure 5d). Multiple RF regressions documented strong culture age and taxonomic effects to Y(II), and a slight fertilizing effect from Fe (Figure 6). However, the exact mechanisms underlying the general inhibiting effects of ash to phytoplankton growth or from the ash 2 treatment (Figure 5cd, Figure 6d,i) are unresolved. While our models did not identify PAH impacts to Y(II) (Figure 6f), PAHs are documented to harm the fluorescence yield, functional absorption cross-section for Photosystem II, and membrane integrity of microalgae (Kottuparambil and Park 2019; Molina and Segura 2021; Othman et al. 2023). We discuss these results and future directions.

Among our samples, the PAH content of air-fall ash was highest, and its values were consistent with reported data. However, due to a lack of available air-fall ash samples, our experimental treatments largely relied on collected and experimentally combusted ash (Table S3). This resulted in reduced the PAH loads in our experimental applications, potentially muting their harmful culture effects. The PAH profiles from air-fall ash were similar to PAHs reported from chaparral and pine forest burns, but 50-80% lower than ash from other ecosystems (Harper et al. 2019). As PAH content is determined by the combusted vegetation (Lima et al. 2005), our results indicate the regions burned during the CZU fire that generated our air-fall ash were composed of similar vegetation taxa present in chaparral and pine stands. Unsurprisingly, the PAH levels of ash collected 2-6 months after the CZU fire decreased, likely due to leaching, volatilization, and because of post-collection drying (Goldfarb 2013). In addition, the fine particle size of our experimentally generated ash indicates a more thorough combustion, which is inconsistent with airborne wildfire ash and lowers PAH content. Future challenge studies that apply wood ash with PAH profiles more aligned with wildfire ash content (e.g., our air-fall ash) may better approximate wildfire impacts.

We assessed the responses to ash in *D. tertiolecta* and *I. galbana*—so-called “bait microalgae”—that are logistically desirable and common in aquaculture and aquarium operations (Creswell 2010; Lombardi and Wangersky 1995; Widowati et al. 2017). Perhaps a result, the maximum Y(II) values observed (0.72-0.76, Figure 5d) show our cultures had near optimal physiological performance. While the Y(II) curves for the control and ash 1 treatments were similar for both species, the ash 2 treatment clearly lost photosynthetic efficiency. Figure 5d shows that for *D. tertiolecta*, Y(II) declined roughly 0.15 and for *I. galbana* Y(II) dropped ∼0.05. Still our phytoplankton cultures appeared robust, attaining elevated Y(II) values that outperformed published assessments of these phytoplankton (Duan et al. 2023; Geider et al. 1998). While we do not understand mechanistically why Y(II) values declined in our ash 2 treatment, wildfire ash exposures in freshwater streams have preceded significant shifts in the community abundance and diversity of microalgae (Earl and Blinn 2003; Klose et al. 2015). Variable responses of marine phytoplankton to ash loads may also drive broad shifts in marine producer communities. Whatever the precise cause for the culture impacts from ash 2, the phytoplankton studied here are not representative taxa of the nearshore Monterey Bay or broader California Current ecosystems (Fischer et al. 2020; Taylor and Landry 2018). To better contextualize the acute impacts of wildfire ash toxicity to regional marine ecosystems, future experiments on taxa common during upwelling regimes (*Chaetoceros, Rhizosolenia, Skeletonema spp*) or in harmful algal blooms (*Pseudo-nitzschia, Akashiwo spp*) may be appropriate models.

This study was opportunistic given our geographic proximity to the CZU fire and the phytoplankton culturing infrastructure at the Monterey Bay Aquarium. Several lessons learned may improve future experimental studies. To begin, our experimental design constrained the structure of several predictor and response variables (Figure 6a-e), which in turn limited the broad utility of the RF model outputs. The strongest variable impact was culture age and species-specific differences (Figure 6f). As this partially reflects the inherent density dependent growth of phytoplankton cultures (Figure 4c), future studies that run longer culture cycles and dose ash on days 8-30 will avoid detecting the initial spike in Y(II) that potentially masked other experimental effects. In addition, future experiments that repeatedly dose cultures (e.g., daily) may more closely mimic wildfire events and provide a sequence of cumulative ash (and associated toxin) values to model against observed Y(II). Such inquiries may better parse the different ability of species and clades of phytoplankton to exhibit resistance (to remain undisturbed) and resilience (to recover after disturbance) to wildfire ash. Broader taxonomic studies may also reveal some microalgae taxa to exhibit extreme sensitivity to ash, to have no resistance and no resilience, and to disappear entirely after exposure.

Undoubtedly, ash applications (Figure 5b) restricted light availability to cultures, inhibiting photosystem II. Experiments using aqueous extracts from ash may avoid ash-induced shading and disaggregate other potential impacts from ash, better isolating the toxicological impacts from wildfire ash to phytoplankton. As we did not measure light availability in cultures, photometry data would also improve understanding baseline culture performance and enhance model utility. (Alternatively, shading impacts from ash could also be tested with chemically inert substances added to cultures.) Further, while weighing filtered culture extracts provided diagnostic insights into culture growth, flow cytometry technology (Marie et al. 2014; Veldhuis and Kraay 2000) would significantly advance experimental possibilities. This might improve growth estimate precision, automate species identification in polycultures, and provide microscopy imagery for additional cellular pathology analysis. Lastly, preparing and maintaining phytoplankton cultures is logistically demanding and requires significant facilities and expertise. As with previous studies (Miller et al. 2021; Nicholson et al. 2020; O’Sullivan et al. 2022), our partnership with a public aquarium was essential.

Our analyses underscore the importance of phytoplankton studies for understanding large scale environmental pollution. As pollutants can concentrate at higher trophic levels, studies of marine top predators have been useful for documenting how anthropogenic pollutants move through and impact ocean ecosystems (Sonne 2010; Storelli et al. 2005; Walsh 2018). However, research documenting pollutant concentrations in producer taxa may be better suited for determining pollutant fluxes (Gagné et al. 2019) and therefore may better facilitate large scale marine pollution monitoring. *Ex situ* studies like the present experiments may be important for establishing the trophic transfer of wildfire pollutants (PAHs, HMs), and present opportunities for testing and controls that field studies cannot provide. Subsequent, complementary field studies may then further establish experimental trophic transfer parameters in wild ecosystems, and reveal the geography of pollutant deposition as well as the relative prevalence of riverine and atmospheric export (Liu et al. 2021).

Beyond deriving global patterns of wildfire pollutants and resolving their ecosystem impacts, a better understanding of the precise physiological mechanisms whereby ash-borne pollutants inhibit phytoplankton communities is needed. HM toxicity, for example, is extensively documented in terrestrial plants (Nagajyoti et al. 2010; Riyazuddin et al. 2021) but is less understood in phytoplankton (Sharma et al. 2021; Zhang and Rickaby 2020). Likewise, PAH toxicity to terrestrial plants is more resolved (Desalme et al. 2013; Molina and Segura 2021) while case studies of phytoplankton impacts are fewer (Kottuparambil and Agusti 2020; Page et al. 2022). A better understanding of both pollutant impacts to phytoplankton—and any differences between beneficial phytoplankton and species forming harmful algal blooms—will help determine the potential significance and scale of future wildfire impacts to marine phytoplankton and ocean ecosystems.

Finally, it is important to set such potential impacts within long-term contexts that include both historical contributions as well as future forecasts arising from climate change. Especially for marine systems, historical studies have provided more informed benchmarks of ocean health that are integral for assessing the current status of ecosystems (Lotze and Worm 2009; McClenachan et al. 2024) and for documenting resiliency and natural climate solutions (Nicholson et al. 2024). Here in this context, historical documentation on the magnitude and occurrence of wildfires (e.g., Keeley and Syphard 2021; Marlon et al. 2012) that directly influence coastal ecosystems is needed. In addition, future projections for forest fires under various climate change regimes (Barbero et al. 2015; Jones et al. 2022) and their potential influence to marine systems are also worthwhile. These analyses must resolve not only physical mechanisms underlying wildfires but the export pathways of terrestrial pollutants and their transport through marine food webs. We hope the size fractionated pollutant content and experimental methods and recommendations that we report here will advance such future work.

## ACKNOWLEDGEMENTS

M. Ezcurra, P. Clarkson, and J. Hoech provided logistical support for the phytoplankton culture experiments at MBA. F. Chavez advised on oceanography, remote sensing and study design. A. Copenhaver, E. Brown, and K. Connor advised on science communication. T. Kana and Bay Instruments loaned technical instrumentation and provided operational training. J. Kiernan, K. Kittleson, and R. Takesue provided in situ wildfire ash samples, J. O’Sullivan provided field images. T. Maddox, R. Poppenga, and C. Perkins performed diagnostic ash analyses and contextualized results. Anonymous reviewers improved earlier versions of the manuscript.

## AUTHOR CONTRIBUTIONS

KV, JL, and CS designed the study. KV, JL, and DV conducted the experiments. AP performed diagnostic analyses. KV performed data analyses, wrote scripts, and generated the figures. KV wrote the manuscript with contributions from JL and AP. All authors reviewed the manuscript.

## DATA AVAILABILITY

Data and code are available at third party repositories, at GitHub (https://bit.ly/3zoUBtr) and at the Open Science Framework (https://osf.io/mk42n/).

## FUNDING

This study was supported by generous contributions from members and donors to the Monterey Bay Aquarium.

## COMPETING INTERESTS

The authors declare no competing interests.

## SUPPLEMENTAL MATERIAL

**Methods**, Phytoplankton culture facilities and preparations

### Tables

Table S1, Metadata for all ash samples and diagnostic testing performed.

Table S2, Tree species and composition of experimentally combusted wood ash. Table S3, Detailed description of the phytoplankton culture treatment regimes.

### Figures

Figure S1, Available 2020 PM_2.5_ time series for 3 measurement stations in Pacific Grove, California.

Figure S2, Separated air-fall wildfire ash and experimentally combusted wood ash.

Figure S3, Other diagnostic PAH ratios and indices for air-fall ash.

## Notes

https://bit.ly/3zoUBtr

https://osf.io/mk42n/

### Competing Interest Statement

The authors have declared no competing interest.

