## Supplemental Material for "Wildfire ash impacts to photosynthesis and growth of marine phytoplankton cultures"

**By Van Houtan et al.**

**FILE CONTENTS:**


**Methods**

Phytoplankton culture facilities and preparations

**Tables**

Table S1, Metadata for all ash samples and diagnostic testing performed.

Table S2, Tree species and composition of experimentally combusted wood ash.

Table S3, Detailed description of the phytoplankton culture treatment regimes.

**Figures**

Figure S1, Available 2020 $PM_{2.5}$ time series for 3 measurement stations in Pacific Grove, California.

Figure S2, Separated air-fall wildfire ash and experimentally combusted wood ash.

Figure S3, Other diagnostic PAH ratios and indices for air-fall ash.

Figure S4, PAH results for wildfire ash samples collected at the $CZU_0$, $CZU_2$, and $CZU_3$ sites.

Figure S5, Repeat of Figure 1e for collected and generated ash samples.

### Methods

*PHYTOPLANKTON CULTURE PREPARATION*

We reared phytoplankton cultures in controlled facilities as part of the regular maintenance and sustainability of bivalve populations (Creswell 2010) that support public-facing exhibits at the Monterey Bay Aquarium. To prepare an experimental array, we first cleaned and sterilized culture containers, using muriatic acid wash (20° Baume HCl, 31.45% concentration, Sunnyside Corporation) to remove organic film and mineral deposits from clear PET carboys (Vintage Shop, 23.3 L). After sterilization, we neutralized the acid-washed carboys with $NaHCO_3$ and rinsed repeatedly with freshwater.

Before adding the phytoplankton cultures, we further prepared and sterilized containers with seawater and aeration. To begin, we added 10ml of household bleach (5.7% NaClO, Pure Bright Germicidal Ultra Bleach) to each carboy. We sourced seawater directly from Monterey Bay (36.6193° N, 121.9006° W), passed it through a 1 μm canister filter (GE Hytrex), and filled each carboy to within 1 cm of the aperture. To add aeration, we inserted a custom-drilled two-hole rubber stopper assembly (#10 solid pure white gum) in the opening of each carboy such that the lower portion of the stopper contacts the seawater-bleach solution, without inducing air bubbles. The two-holed rubber stopper assembly was pre-fitted with a ~50 cm long elbow glass tube (out diameter 7mm, custom-made) that extends to nearly the bottom of the carboy, serving as the air inlet. A second short glass tube elbow (~5 cm long) that just clears the bottom of the stopper serves as the air outlet. These prepared containers cured for 12 hours.

To prepare for inoculation, we used a cleaned, clear PVC siphon hose (25 mm OD, 19 mm ID, 2.44 m length) to remove ~2 L of seawater from each carboy until the water level reaches the maximum inner chamber diameter. To neutralize the bleach, we added 5 ml of sodium thiosulphate solution (159 g of $Na_2S_2O_3$ pentahydrate crystals, HP grade, Gallade Chemical, Inc., dissolved in 1 L diH2O) to each container. We then connected the inlet tubing of each carboy to an air source with an intermediate 50 mm HEPA-vent in-line disk filter (Hepa-Vent 0.3 μm, Whatman™), allowing the contents to aerate at a moderate rate for 1 hour.

| DATE | AGE (months) | ASH TYPE | SAMPLE ID | LOCATION | COLLECTOR | LAT (° N) | LON (° W) | PAH | HM | SIA |
|---|---|---|---|---|---|---|---|---|---|---|
| 8/20/20 | 0.0 | air-fall | deposited | Pacific Grove, CA | MBA/Van Houtan | 36.618138 | -121.906306 | 6 | 4 | -- |
| 11/5/20 | 2.5 | collected | $CZU_0$ | CZU complex | NOAA/Kiernan | 37.076411 | -122.219378 | 3 | -- | -- |
| 12/12/20 | 3.7 | collected | $CZU_1$ | CZU complex | MBA/Van Houtan | 37.104821 | -122.143461 | 18 | 18 | 18 |
| 3/5/21 | 6.5 | collected | $CZU_2$ | CZU complex | SC/Kittleson | 37.072905 | -122.230569 | 6 | -- | -- |
| 3/6/21 | 6.5 | collected | $CZU_3$ | CZU complex | USGS/Takesue | 37.06209 | -122.22861 | 6 | -- | -- |
| 2/15/21 | 0.0 | generated | burn | Pacific Grove, CA | MBA/Van Houtan | n/a | n/a | 18 | 20 | -- |

**Table S1. Metadata for all ash samples and diagnostic testing performed.** "Date" is collection date, "Age (months)" is the estimated time in months elapsed between the burn and collection date, and "Sample ID" represents internal identifiers. Here, "PAH", "HM", and "SIA" are the number of samples analyzed for polycyclic aromatic hydrocarbons ($n$=57), heavy metals ($n$=42), and stable isotope ($n$=18), respectively, where the analyses of sieved and size fractionated samples are underlined. Figure S2 provided further details for collected samples at the CZU sites.

| FAMILY | GENUS | SPECIES | COMMON | PARTS | PROPORTION |
|---|---|---|---|---|---|
| Cupressaceae | Sequoia | sempervirens | Coastal redwood | foliage, twigs, branches | 0.5 |
| Fagaceae | Quercus | agrifolia | California Live Oak | foliage, twigs, branches | 0.2 |
| Pinaceae | Pinus | radiata | Monterey Pine | foliage, twigs, branches | 0.1 |
| Myrtaceae | Eucalyptus | globulus | Tasmanian blue gum | foliage, twigs, bark strips | 0.05 |
| Myrtaceae | Eucalyptus | camaldulensis | River red gum | foliage, twigs | 0.05 |
| Cupressaceae | Cupressus | macrocarpa | Monterey Cypress | foliage, branches | 0.05 |
| Pinaceae | Abies | balsamea | Balsam Fir | foliage, twigs | 0.05 |

**Table S2. Tree species composition for the experimentally combusted wood ash.** .
Physical and chemical profile of this combusted ash is plotted in Figure 3 of the main text.

| Treatment | Carboy | Phytoplankton | Ash (g) | Airfall % | $CZU_0$ % | $CZU_1$ % | $CZU_2$ % | $CZU_3$ % | Generated % |
|---|---|---|---|---|---|---|---|---|---|
| postitive control | CO1 | *D. teriolecta* | 0.0 | 0 | 0 | 0 | 0 | 0 | 0 |
| postitive control | CO2 | *I. galbana* | 0.0 | 0 | 0 | 0 | 0 | 0 | 0 |
| negative control | NC1 | -- | 28.4 | 0.0 | 0.0 | 52.9 | 31.7 | 15.3 | 0.0 |
| negative control | NC2 | -- | 23.4 | 0.0 | 0.0 | 70.1 | 15.3 | 14.6 | 0.0 |
| ash 1 | DA1 | *D. tertiolecta* | 28.5 | 3.3 | 88.3 | 8.4 | 0.0 | 0.0 | 0.0 |
| ash 1 | DA2 | *I. galbana* | 28.3 | 3.3 | 88.6 | 8.1 | 0.0 | 0.0 | 0.0 |
| ash 2 | CA1 | *D. teriolecta* | 28.3 | 0.0 | 0.0 | 55.3 | 0.0 | 0.0 | 44.7 |
| ash 2 | CA2 | *I. galbana* | 28.3 | 0.0 | 0.0 | 55.0 | 0.0 | 0.0 | 45.0 |

**Table S3. Experimental treatment, ash dosing, and control regimes for the phytoplankton culture study.** Full details of the experimental manipulations of the phytoplankton cultures testing the potential effects of combusted wood ash on photosynthesis and growth. “Treatment” is the experimental regime, “Carboy” is the culture container label, “Phytoplankton” is the cultured species, and “Ash (g)” is the total ash added in g. We added an average of 28.35 g of ash to the 23L of phytoplankton culture (1.23 g $L^{-1}$) in each carboy. The remaining columns are the percentages of the various ash types comprising the “Ash (g)” entry. Based on the density dependent growth pattern in Figure 3, each culture was run for 12 days. Due to logistical constraints, each trial was only replicated for each species. Phytoplankton value denoted by “—” or ash mass “0” means nothing added, describing the negative and positive controls, respectively.

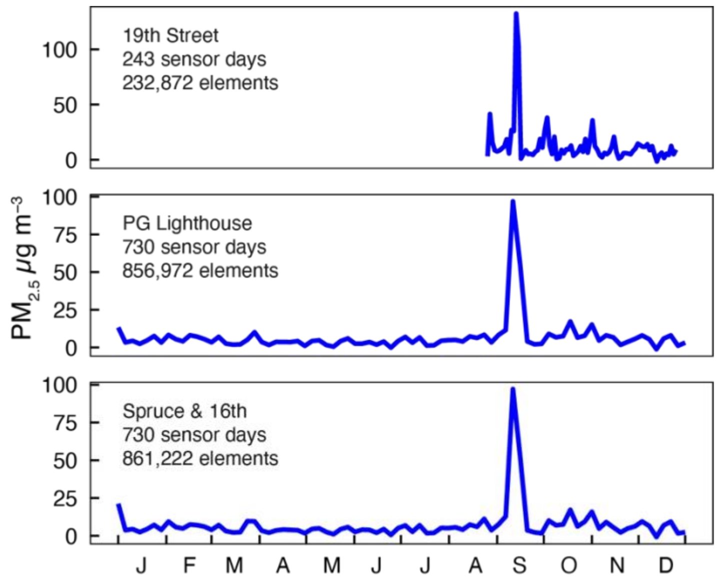


**Figure S1. Full comparison of individual time series of all Purple Air station sensors modeled in Figure 1a.** Three sensors used were the closest available for the 2020 calendar year, from the publicly available community science map portal provided by Purple Air at https://bit.ly/3GsdpMw. Station name, days active in 2020, and total number of instrument readings is listed in each panel. Each station contains 2 sensor channels (A, B) which for these 3 stations together provided 1,703 total sensor days and 1,951,066 measurements in 2020. The EPA uses the $PM_{2.5}$ metric in their reported Air Quality Index, considering values > 50 µg $m^{-3}$ hazardous to human health.

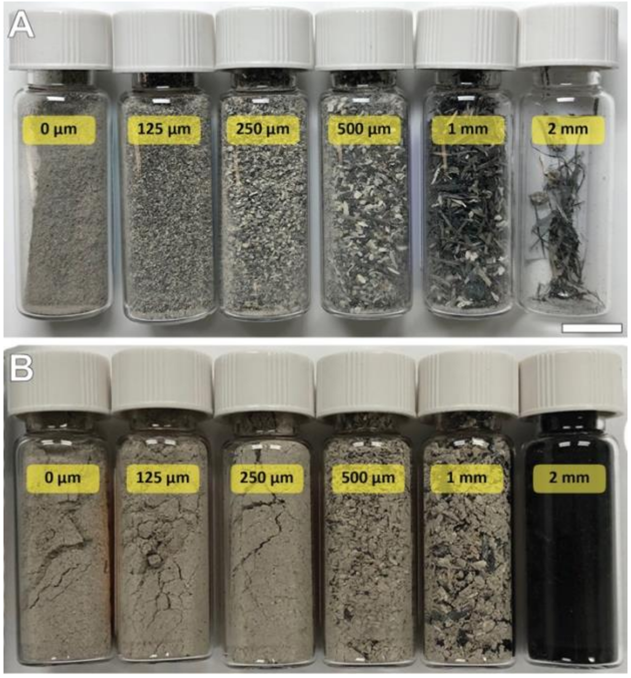


**Figure S2. Size-fractionated wood ash in glass vials.** (A) Air-fall wildfire ash collected in pre-cleaned glassware on the Monterey Bay Aquarium roof on 20-21 August 2020 during the CZU Lightning Complex fires. (B) Experimentally generated ash combusted from collected tree parts (see Table S2). Measurements listed on the vials refers to the sieve pore size on which the samples were retained. All ash particles were separated with US standard (ASTM E-11) stainless steel sieves. Figures 1-2 in the main text plot the size composition of each ash type.

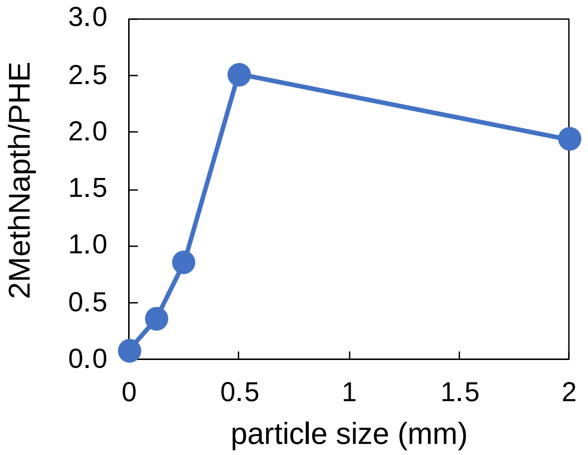


**Figure S3. Other diagnostic PAH ratios and indices for air-fall ash.** The 2 PAH ratios plotted in Figure 1f, FLA/(FLA+PYR) and ANT/(ANT+PHE), are considered diagnostic in determining the pollutant origin. Here we provide additional results. This panel describes the 2MethNapth/PHE index, which is the ratio of 2-methyl naphthalene to phenanthrene. Due to undetermined PAH quantities for 1mm particle samples, there are no 2MethNapth/PHE results for this size class. In addition to these size fractionated data, the whole sample index value for 2MethNapth/PHE = 1.07. For the 2MethNapth/PHE ratio, values < 1 indicate biomass combustion, and values ranging from 2-6 indicate fossil fuel combustion (Takesue et al 2023). Both the 500μm and 2mm particle samples are > 2, indicating a fossil fuel origin for these air-fall samples. This putative determination is contradicted by a visual examination of the 2mm particles that appear to be partially combusted conifer needles (e.g., Figure 1b, Figure S3a). Despite undetermined quantities for all particle size classes except 125 μm ash particles, here we also report (not plotted here) on 2 additional PAH indices. IP/(IP+BghiP) is an index of indeno(1,2,3-cd)pyrene and benzo(g,h,i)perylene and BaP/BghiP is an index of benzo(a)pyrene and benzo(g,h,i)perylene. For air-fall ash particles 125 μm in size, the IP/(IP+BghiP) = 0.50, a borderline value between a petroleum combustion source and biomass combustion. Also for air-fall ash particles 125 μm in size, the BaP/BghiP = 0.56, indicating the combustion source did not originate from automobile traffic emissions.
